# Domain-specific working memory, but not dopamine-related genetic variability, shapes reward-based motor learning

**DOI:** 10.1101/524900

**Authors:** Peter Holland, Olivier Codol, Elizabeth Oxley, Madison Taylor, Elizabeth Hamshere, Shadiq Joseph, Laura Huffer, Joseph M. Galea

## Abstract

The addition of rewarding feedback to motor learning tasks has been shown to increase the retention of learning, spurring interest in the possible utility for rehabilitation. However, laboratory-based motor tasks employing rewarding feedback have repeatedly been shown to lead to great inter-individual variability in performance. Understanding the causes of such variability is vital for maximising the potential benefits of reward-based motor learning. Thus, using a large cohort (n=241) we examined whether spatial (SWM), verbal (VWM) and mental rotation (RWM) working memory capacity and dopamine-related genetic profiles were associated with performance in two reward-based motor tasks. The first task assessed participant’s ability to follow a hidden and slowly shifting reward region based on hit/miss (binary) feedback. The second task investigated participant’s capacity to preserve performance with binary feedback after adapting to the rotation with full visual feedback. Our results demonstrate that higher SWM is associated with greater success and a greater capacity to reproduce a successful motor action, measured as change in reach angle following reward. Whereas higher RWM was predictive of an increased propensity to express an explicit strategy when required to make large adjustments in reach angle. Therefore, both SWM and RWM were reliable predictors of success during reward-based motor learning. Change in reach direction following failure was also a strong predictor of success rate, although we observed no consistent relationship with any type of working memory. Surprisingly, no dopamine-related genotypes predicted performance. Therefore, working memory capacity plays a pivotal role in determining individual ability in reward-based motor learning.

**Significance statement:** Reward-based motor learning tasks have repeatedly been shown to lead to idiosyncratic behaviours that cause varying degrees of task success. Yet, the factors determining an individual’s capacity to use reward-based feedback are unclear. Here, we assessed a wide range of possible candidate predictors, and demonstrate that domain-specific working memory plays an essential role in determining individual capacity to use reward-based feedback. Surprisingly, genetic variations in dopamine availability were not found to play a role. This is in stark contrast with seminal work in the reinforcement and decision-making literature, which show strong and replicated effects of the same dopaminergic genes in decision-making. Therefore, our results provide novel insights into reward-based motor learning, highlighting a key role for domain-specific working memory capacity.

## Introduction

When performing motor tasks under altered environmental conditions, adaptation to the new constraints occurs through the recruitment of a variety of systems (Taylor and Ivry, 2014). Arguably the most studied of those systems is cerebellum-dependent adaptation, which consists of the implicit and automatic recalibration of mappings between actual and expected outcomes, through sensory prediction errors (Morehead et al., 2017; Tseng et al., 2007). Besides cerebellar adaptation, other work has demonstrated the involvement of a more cognitive, deliberative process whereby motor plans are adjusted based on an individual’s structural understanding of the task (Bond and Taylor, 2015; Taylor and Ivry, 2011). We label this process ‘explicit control’ (Codol et al., 2018; Holland et al., 2018) but it has also been referred to as strategy (Taylor and Ivry, 2011) or explicit re-aiming (Morehead et al., 2015). Recently it has been proposed that reinforcement learning, whereby the memory of successful or unsuccessful actions are strengthened or weakened, respectively, may also play a role (Huang et al., 2011; Izawa and Shadmehr, 2011; Shmuelof et al., 2012). Such reward-based reinforcement has been assumed to be an implicit and automatic process (Haith and Krakauer, 2013). However, recent evidence suggests that phenomena attributed to reinforcement-based learning during visuomotor rotation tasks can largely be explained through explicit processes (Codol et al., 2018; Holland et al., 2018).

One outstanding feature of reinforcement-based motor learning is the great variability expressed across individuals (Codol et al., 2018; Holland et al., 2018; Therrien et al., 2016, 2018). What factors underlie such variability is unclear. If reinforcement is indeed explicitly grounded, it could be argued that individual working memory capacity (WMC), which reliably predicts propensity to employ explicit control in classical motor adaptation tasks (Anguera et al., 2010; Christou et al., 2016; Sidarta et al., 2018), would also predict performance in a reinforcement-based motor learning task. If so, this would strengthen the proposal that reward based motor learning bears a strong explicit component. Anguera et al., (2010) demonstrated that mental rotation WM (RWM), unlike other forms of working memory such as verbal working memory (VWM), correlates with explicit control. More recently, Christou et al. (2016) reported a similar correlation with spatial WM (SWM).

Another potential contributor to this variability is genetic profile. In previous work (Codol et al., 2018; Holland et al., 2018), we argue that reinforcement-based motor learning performance relies on a balance between exploration and exploitation of the task space, a feature reminiscent of structural learning and reinforcement-based decision-making (Daw et al., 2005; Frank et al., 2009; Sutton and Barto, 1998). A series of studies from Frank and colleagues suggests that individual tendencies to express explorative/exploitative behaviour can be predicted based on dopamine-related genetic profile (Doll et al., 2016; Frank et al., 2007, 2009). Reinforcement has consistently been linked to dopaminergic function in a variety of paradigms, and thus, such a relationship could also be expected in reward-based motor learning (Pekny et al., 2015). Specifically, Frank and colleagues focused on Catecholamine-O-Methyl-Transferase (COMT), Dopamine- and cAMP-Regulated neuronal Phosphoprotein (DARPP32) and Dopamine Receptor D2 (DRD2), and suggest a distinction between COMT-modulated exploration and DARPP32- and DRD2-modulated exploitation (Frank et al., 2009).

Consequently, we investigated the influence of WM capacity (RWM, SWM, and VWM) and genetic variations in dopamine metabolism (DRD2, DARP32, and COMT) on an individual’s ability to perform reward-based motor learning. We examined this using two established reward-based motor learning tasks. First, a task analogous to a gradually introduced rotation (Holland et al., 2018) required participants to learn to adjust the angle at which they reached to a slowly and secretly shifting reward region (Acquire); second, a task with an abruptly introduced rotation (Codol et al., 2018; Shmuelof et al., 2012) required participants to preserve performance with reward-based feedback after adapting to a visuomotor rotation (Preserve). The use of these two tasks enabled us to examine whether similar predictors of performance explained participant’s capacity to acquire and preserve behaviour with reward-based feedback.

## Methods

Prior to the start of data collection, the sample size, variables of interest and analysis method were pre-registered. The pre-registered information, data and analysis code can be found online at https://osf.io/j5v2s/ and https://osf.io/rmwc2/ for the Preserve and Acquire tasks, respectively.

### Participants

121 (30 male, mean age: 21.06, range: 18-32) and 120 (16 male, mean age: 19.24, range: 18-32) participants were recruited for the Acquire and Preserve tasks, respectively. All participants provided informed consent and were remunerated with either course credit or money (£7.50/hour). All participants were free of psychological, cognitive, motor or uncorrected visual impairment. The study was approved by and performed in accordance with the local research ethics committee of the University of Birmingham, UK.

### Experimental design

Participants were seated before a horizontally fixed mirror reflecting a screen placed above, on which visual stimuli were presented. This arrangement resulted in the stimuli appearing at the level on which participants performed their reaching movements. The Acquire (gradual) and Preserve (abrupt) tasks were performed on two different stations, with a KINARM (BKIN Technology, London, Ontario; sampling rate: 1000Hz) and a Polhemus 3SPACE Fastrak tracking device (Colchester, Vermont U.S.A.; sampling rate: 120Hz), employed respectively. The Acquire task was run using Simulink (The Mathworks, Natwick, MA) and Dexterit-E (BKIN Technology), while the Preserve task was run using Matlab (The Mathworks, Natwick, MA) and Psychophysics toolbox (Brainard, 1997). The Acquire task employed the same paradigm and equipment as Holland et al. (2018), with the exception of the maximum reaction time (RT) which was increased from 0.6s to 1s, and the maximum movement time (MT) which was reduced from 1s to 0.6s. The Preserve task used the same setup and display as in Codol et al. (2018); however, the number of ‘refresher’ trials during the binary feedback (BF) blocks was increased from one to two in every 10 trials. The designs were kept as close as possible to their respective original publications to promote replication and comparability across studies. In both tasks reaching movements were made with the dominant arm.

### Reaching tasks

#### Acquire task

Participants performed 670 trials, each of which followed a stereotyped timeline. The starting position for each trial was in a consistent position roughly 30cm in front of the midline and was indicated by a red circle (1cm radius). After holding the position of the handle within the starting position, a target (red circle, 1cm radius) appeared directly in front of the starting position at a distance of 10cm. Participants were instructed to make a rapid ‘shooting’ movement that passed through the target. If the cursor position at a radial distance of 10cm was within a reward region (±5.67°, initially centred on the visible target; grey region in Figure 1a) the target changed colour from red to green and a green tick was displayed just above the target position, informing participants of the success of their movement. If, however, the cursor did not pass through the reward region, the target disappeared from view and no tick was displayed, signalling failure (binary feedback). After each movement, the robot returned to the starting position and participants were instructed to passively allow this.

**Figure 1:**
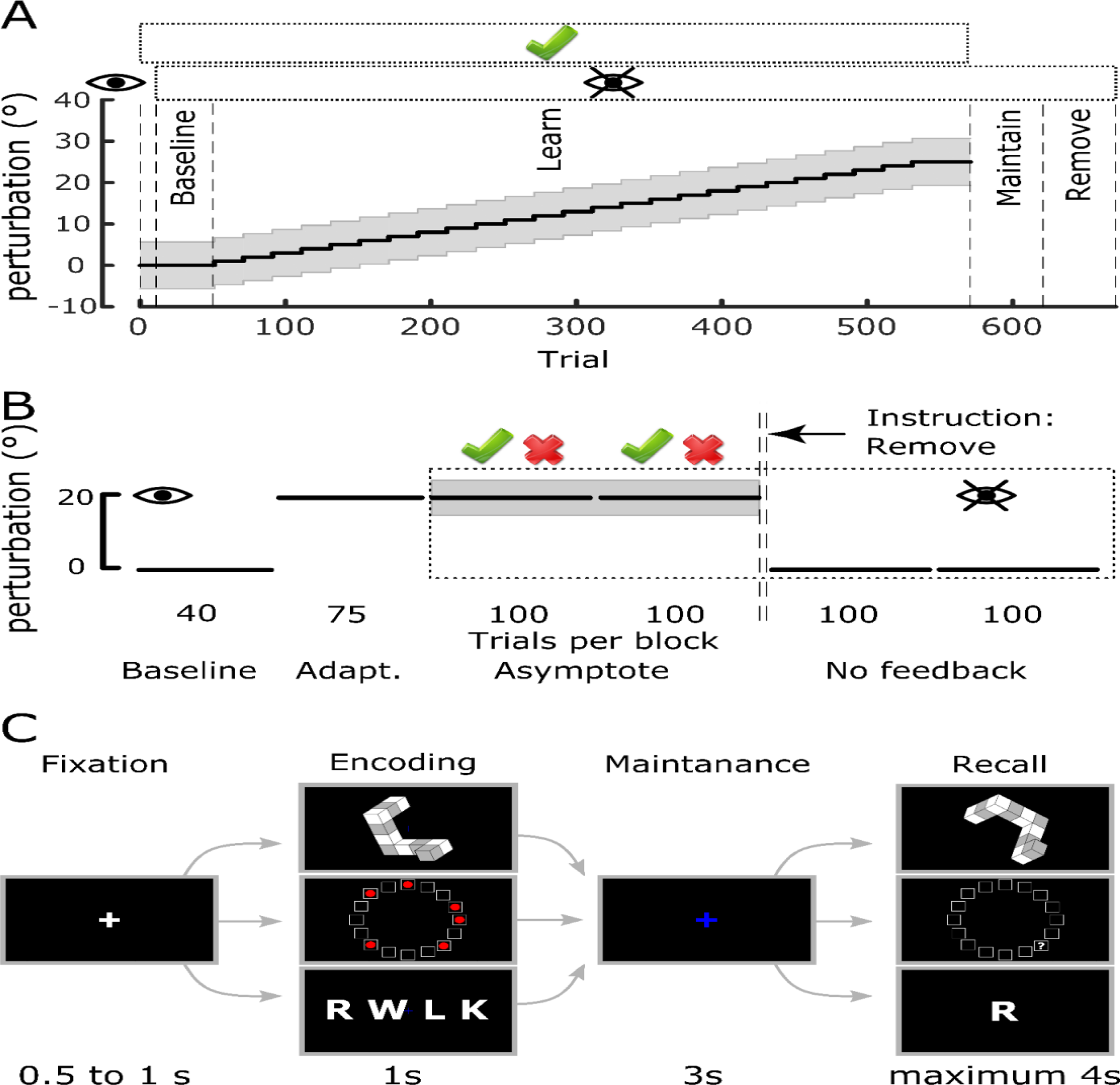
Experimental design. **A:** Time course of the Acquire task with the different experimental periods labelled. The grey region represents the reward region, which gradually rotated away from the visual target after the initial baseline period. The rectangle enclosing the green tick above the axes represents trials in which reward was available, and the rectangle with the ‘eye’ symbol indicates when vision was not available. **B:** Time course of the Preserve task. After adapting to an initial rotation with vision available, vison was removed (eye symbol) and reward-based feedback was introduced (tick and cross above the axes). Prior to the no-feedback blocks participants were instructed to remove any strategy they had been using. **C:** WM capacity tasks, the three tasks followed a stereotyped timeline with only the items to be remembered differing. Each trial consisted of 4 phases (Fixation, Encoding, Maintenance, and Recall) with the time allocated to each displayed below.

For the first 10 trials, the position of the robotic handle was displayed as a white cursor (0.5 cm radius) on screen, following this practice block the cursor was extinguished for the remainder of the experiment and participants only received binary feedback. The baseline block consisted of the first 40 trials under binary feedback, during this period the reward region remained centred on the visible target. Subsequently, unbeknownst to the participant the reward region rotated in steps of 1° every 20 trials; the direction of rotation was counterbalanced across participants. After reaching a rotation of 25° the reward region was held constant for an additional 20 trials. Performance during these last 20 trials was used to determine overall task success. Subsequently, binary feedback was removed, and participants were instructed to continue reaching as they were (maintain block) for the following 50 trials. Following this, participants were then informed that the reward region shifted during the experiment but not of the magnitude or the direction of the shift. They were then instructed to return to reaching in the same manner as they were at the start of the experiment (remove block, 50 trials). During the learning phase of the task participants were given a 1-minute rest after trials 190 and 340.

#### Preserve task

Participants performed 515 trials in total. On each trial participants were instructed to make a rapid ‘shooting’ movement that passed through a target (white circle, radius: 0.125cm) visible on the screen. The starting position for each trial was indicated by a white square (width: 1cm) roughly 30cm in front of the midline and the target was located at angle of 45⁰ from the perpendicular in a counter clockwise direction at a distance of 8cm. The position of the tracking device attached to the fingertip was displayed as a cursor (green circle, radius: 0.125cm). When the radial distance of the cursor from the starting position exceeded 8cm, the cursor feedback disappeared, and the end position was displayed instead. First, participants performed a baseline period of 40 trials, during which the position of the cursor was visible and the cursor accurately reflected the position of the fingertip. In the adaptation block (75 trials), participants were exposed to an abruptly introduced 20° clockwise visuomotor rotation of the cursor feedback (Figure 1b). Subsequently, all visual feedback of the cursor was removed, and participants received only binary feedback. If the end position of the movement fell within a reward region, the trial was considered successful and a tick was displayed; otherwise a cross was displayed. The reward region was centred at a clockwise rotation of 20° with respect to the visual target with a width of 10° i.e. it was centred on the direction that successfully accounted for the previously experienced visuomotor rotation. Binary feedback was provided for 200 trials divided into 2 blocks of 100 trials (asymptote blocks). Furthermore, participants experienced 2 “refresher” trials for every 10 trials, where rotated visual feedback of the cursor position was again accessible (Codol et al., 2018; Shmuelof et al., 2012). This represents an increase compared to Codol et al. (2018) because participants in this study tended to have poorer performance under binary feedback, possibly due to a fatigue effect following the WM tasks (Anguera et al., 2012; see discussion).

Finally, two blocks (100 trials each) with no performance feedback were employed in order to assess retention of the perturbation (no-feedback blocks). Before the first of those two blocks, participants were informed of the visuomotor rotation, asked to stop accounting for it through aiming off target and to aim straight at the target.

### Working memory tasks

Participants performed three WM tasks, all of which followed the same design with the exception of the nature of the items to be remembered (Figure 1c). All WM tasks were run using Matlab (The Mathworks, Natwick, MA) and Psychophysics toolbox (Brainard, 1997). At the start of each trial, a white fixation cross was displayed in the centre of the screen for a period of 0.5 to 1s randomly generated from a uniform distribution (fixation period Figure 1c). In the encoding period, the stimuli to be remembered was displayed for 1s and then subsequently replaced with a blue fixation cross for the maintenance period which persisted for 3s. Finally, during the recall period, participants were given a maximum of 4s to respond by pressing one of three keys on a keyboard with their dominant hand. The ‘1’ key indicated that the stimuli presented in the recall period was a ‘match’ to that presented in the encoding period, the ‘2’ key indicated a ‘non-match’ and pressing ‘3’ indicated that the participant was unsure as to the correct answer. Each WM task contained 5 levels of difficulty with the 12 trials presented for each; 6 of which were trials in which ‘match’ was the correct answer and 6 in which ‘non-match’ was the correct answer. Consequently, each task consisted of 60 trials and the order in which the tasks were performed was pseudorandomised across participants. Prior to the start of each task participants performed 10 practice trials to familiarise themselves with the task and instructions. For both the Acquire and Preserve tasks, the WM tasks were performed in the same experimental session as the reaching. However, in the case of the Acquire task the WM tasks were performed after the reaching task whereas for the Preserve task the WM tasks were performed first.

In the rotation WM task (RWM, Figure 1c top row), the stimuli consisted of six 2D representations of 3D shapes drawn from an electronic library of the Shepard and Metzler type stimuli (Peters and Battista, 2008). The shape presented in the recall period was always the same 3D shape presented in the encoding period after undergoing a screen-plane rotation of 60°, 120°, 180°, 240° or 300°. In ‘match’ trials, the only transform applied was the rotation; however, in ‘non-match’ trials an additional vertical-axis mirroring was also applied. The difficulty of mental rotation has been demonstrated to increase with larger angles of rotation (Shepard and Metzler, 1971) and therefore the different degrees of rotation corresponded to the 5 levels of difficulty. However, given the symmetry of two pairs of rotations (60 and 300, 120 and 240), these 5 levels were collapsed to 3 for analysis.

In the spatial WM task (SWM, Figure 1c middle row), stimuli in the encoding period consisted of a variable number of red circles placed within 16 squares arranged in a circular array (McNab and Klingberg, 2008). In the recall period, the array of squares was presented without the red circles and instead a question mark appeared in one of the squares. Participants then answered to the question “*Was there a red dot in the square marked by a question mark?*” by pressing a corresponding key. In ‘match’ trials the question mark appeared in one of the squares previously containing a red circle and in ‘non-match’ trials it appeared in a square that was previously empty. Difficulty was scaled by varying the number of red circles (i.e. the number of locations to remember) from 3 to 7.

In the verbal WM task (VWM, Figure 1c bottom row), participants were presented with a list of a variable number of consonants during the encoding period. In the recall period a single consonant was presented, and participants answered to the question “*Was this letter included in the previous array?*”. Thus, the letter could either be drawn from the previous list (‘match’ trials) or have been absent from the previous list (‘non-match’ trials). Difficulty in this task was determined by the length of the list to be remembered, ranging from 5 to 9.

### Genetic sample collection and profiling

COMT is thought to affect DA function mainly in the prefrontal cortex (Egan et al., 2001; Goldberg et al., 2003), a region known for its involvement in WM and strategic planning (Anguera et al., 2010; Doll et al., 2015), whereas DARPP32 and DRD2 act mainly in the basal ganglia to promote exploitative behaviour, possibly by promoting selection of the action to be performed (Frank et al., 2009). Consequently, we focused here on SNPs related to those genes: RS4680 (COMT) and RS907094 (DARPP32). Regarding DRD2, there are two potential SNPs available, RS6277 and RS1800497. Although previous studies focusing on exploration and exploitation have assessed RS6277 expression (Doll et al., 2016; Frank et al., 2007, 2009), it should be noted that this SNP varies greatly across ethnic groups, with some allelic variations being nearly completely absent in non-Caucasian-European groups (e.g. see RS6277 in 1000 Genomes Project (The 1000 Genomes Project Consortium et al., 2015)). This has likely been inconsequential in previous work, as Caucasian-European individual represented the majority of sampled groups; here however, this represents a critical shortcoming, as we aim at investigating a larger and more representative population including other ethnic groups. Consequently, we based our analysis on the RS1800497 allele of the DRD2 gene (Pearson-Fuhrhop et al., 2013).

At the end of the task, participants were asked to produce a saliva sample which was collected, stabilized and transported using Oragene.DNA saliva collection kits (OG-500, DNAgenotek, Ontario, Canada). Participants were requested not to eat or drink anything except water for at least two hours before sample collection. Once data collection was completed across all participants, the saliva samples were sent to LGC (Hoddeson, Hertfordshire; https://www.lgcgroup.com/) for DNA extraction (per Oragene protocols: https://www.dnagenotek.com/) and genotyping. SNP genotyping was performed using the KASP SNP genotyping system. KASP is a competitive allele-specific PCR incorporating a FRET quencher cassette. Specifically, the SNP-specific KASP assay mix (containing two different, allele specific, competing forward primers) and the universal KASP master mix (containing FRET cassette plus Taq polymerase in an optimised buffer solution) were added to DNA samples and a thermal cycling reaction performed, followed by an end-point fluorescent read according to the manufacturer’s protocol. All assays were tested on in-house validation DNA prior to being run on project samples. No-template controls were used, and 5% of the samples had duplicates included on each plate to enable the detection of contamination or non-specific amplification. All assays had over 90% call rates. Following completion of the PCR, all genotyping reaction plates were read on a BMG PHERAStar plate reader. The plates were recycled until a laboratory operator was satisfied that the PCR reaction had reached its endpoint. In-house Kraken software then automatically called the genotypes for each sample, with these results being confirmed independently by two laboratory operators. Furthermore, the duplicate saliva samples collected from 5% of participants were checked for consistency with the primary sample. No discrepancies between primary samples and duplicates were discovered.

### Data analysis

#### Acquire task

Reach trials containing MTs over 0.6s or less than 0.2s were removed from analysis (6.9% of trials). The end point angle of each movement was defined at the time when the radial distance of the cursor exceeded 10cm. This angle was defined in relation to the visible target with positive angles indicating clockwise rotations, end point angles and target angles for participants who experienced the counter clockwise rotations were sign-transformed. The explicit component of retention was defined as the difference between the mean reach angle of the maintain block and the remove block, while the implicit component was the difference between the mean reach angle of the remove block and baseline. If during the final 20 trials before the maintain block a participant achieved a mean reach angle within the reward region, participants were considered “*successful*” in learning the rotation; they were considered “*unsuccessful*” otherwise. For regression analysis a binary variable “*task success*” was defined as 1 and 0 for successful and unsuccessful participants, respectively. As in Holland et al (2018), for unsuccessful participants, the largest angle of rotation at which the mean reach angle fell within the reward region was taken as the end of successful performance, and only trials prior to this point were included for further analysis. Success rate was defined as the percentage of trials during the learning blocks in which the end point angle was within the reward region. In order to examine the effect of reward on the change in end point angle on the subsequent trial, we calculated the absolute change in end point angle between consecutive trials (Holland et al., 2018; Sidarta et al., 2018; Therrien et al., 2016, 2018). Subsequently we calculated the median absolute change in angle following rewarded trials (ΔR) and the median absolute deviation of these values (MAD [ΔR]). This analysis was repeated for the changes in angle following unsuccessful trials (ΔP and MAD [ΔP]).

#### Preserve task

Reach trials containing MTs over 1s were removed from analysis (2.38% of trials). The end point angle for each movement was defined at the time that the radial distance of the cursor from the start position exceeded 8cm. Trials in which the error was greater than 80° were excluded from further analysis (0.94% of trials). For each participant we plotted the reach error in one trial against the change in reach angle in the following trial for all trials in the adaptation block. The angle of the line of best fit was then used as the learning rate (Hutter and Taylor, 2018). Using this approach, a perfect adaptation leads to a value of −1, indicating that the error on a given trial is fully accounted for on the next trial. Overall this approach fitted the data well (mean R^2^=0.5038, SD=0.12). As in Codol et al (2018), success rate, corresponding to percentage of rewarded trials, was measured separately in the first 30 and last 170 trials of the asymptote blocks and labelled early and late success rate, respectively. This reflects a dichotomy between a dominantly exploration-driven early phase and a later exploitation-driven phase. Implicit retention was defined as the difference between the average baseline reach direction and the mean reach direction of the last 20 trials of the last no-feedback block (Codol et al., 2018). Analysis of changes in reach angle following rewarded trials were not pre-registered, but were included post-hoc.

#### Exploratory analysis of reaching data

In order to understand which outcome variables in the reaching tasks were predictive of overall task success, we split the learning period into two sections for every participant. We assessed trial-by-trial changes in end point angle in the first section, and compared them to success rate in the second section. For the Acquire task, we assessed trial-by-trial adjustments during the learning block, excluding the final 20 trials, and compared them to success rate in the last 20 trials of the learning block. In the Preserve task, we measured adjustments in the first 100 trials of the asymptote blocks and compared them to success rate in the last 100 trials of the asymptote blocks.

Two additional post-hoc stepwise regressions were performed on data from the Preserve task with early and late success rate the outcome variables and the same set of seven predictors (see statistical analysis section), however, in this case only data from participants with a success rate of greater than 40% was included (N=70).

#### WM tasks

WM performance was defined as the average percentage of correct responses across the 3 highest levels of difficulty for each task. In the case of the RWM task, the symmetrical arrangement of the angles of rotation in effect produced three levels of difficulty and therefore all trials were analysed.

#### Genetics

Genes were linearly encoded, with heterozygote alleles being 0, homozygote alleles bearing the highest dopaminergic state being 1, and homozygote alleles bearing the lowest dopaminergic state being −1 (Table 1). All groups were assessed for violations of the Hardy-Weinberg equilibrium. The participant pool in the Preserve task was in Hardy-Weinberg equilibrium for all three genes considered, even when restricted to the Caucasian-only subpopulation. In the Acquire task population, COMT and DRD2 were in Hardy-Weinberg equilibrium, but DARPP32 was not (p=0.002), with too few heterozygotes. Therefore, the DARPP32 alleles were recoded, with the heterozygotes (0) and the smallest homozygote group (C:C, −1) combined and recoded as 0. In the analysis including only the Caucasian subset, all three alleles were in the Hardy-Weinberg equilibrium, although combining the heterozygote and smallest homozygote group did not change the results.

**Table 1:**
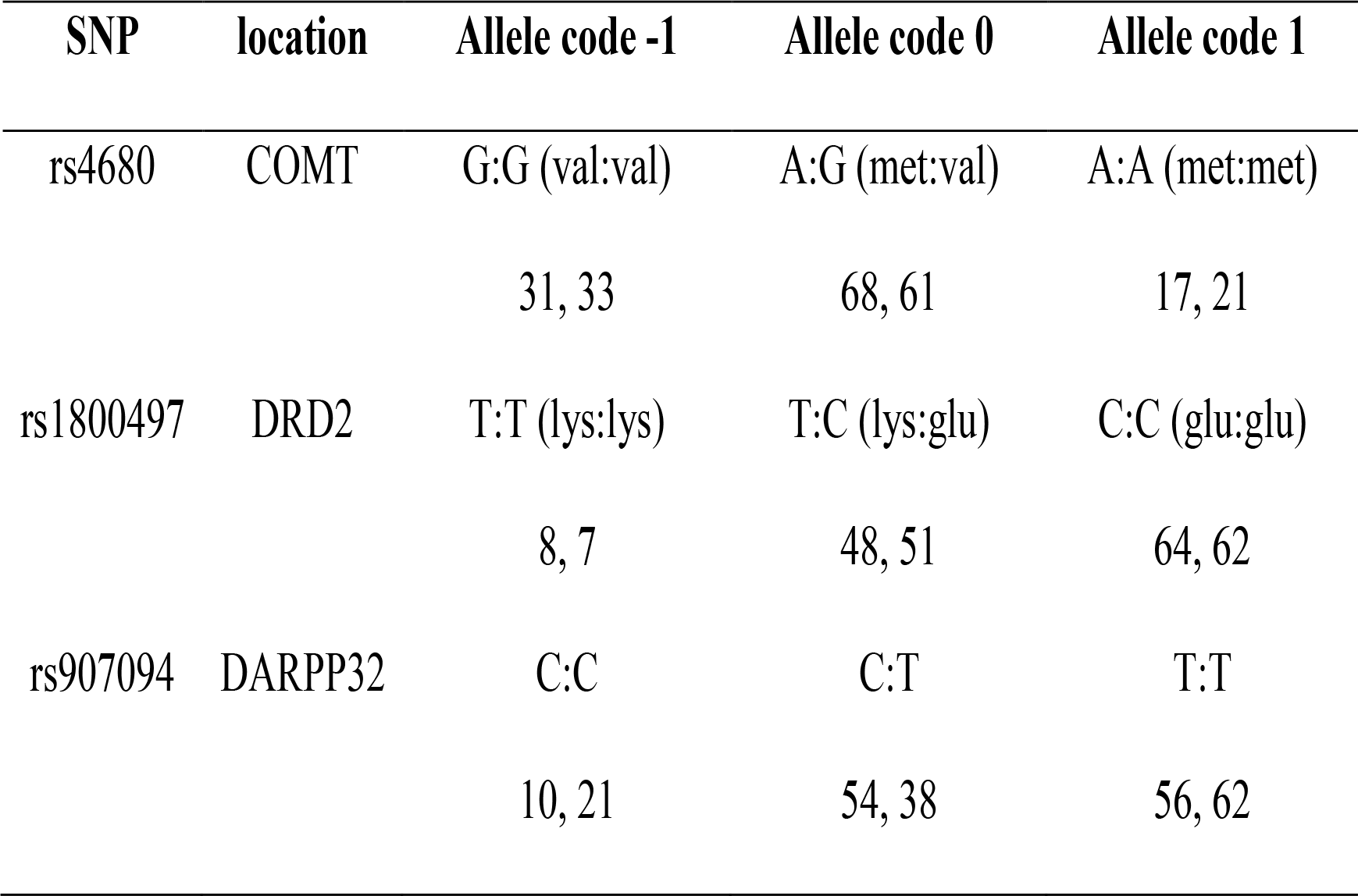
Coding for SNPs. The name of the SNP is provided along with the code assigned to each allele. The numbers represent the counts for the specific allele in the two tasks (Preserve, Acquire).

| SNP | location | Allele code -1 | Allele code 0 | Allele code 1 |
| --- | --- | --- | --- | --- |
| rs4680 | COMT | G:G (val:val) | A:G (met:val) | A:A (met:met) |
|  |  | 31, 33 | 68, 61 | 17, 21 |
| rs1800497 | DRD2 | T:T (lys:lys) | T:C (lys:glu) | C:C (glu:glu) |
|  |  | 8, 7 | 48, 51 | 64, 62 |
| rs907094 | DARPP32 | C:C | C:T | T:T |
|  |  | 10, 21 | 54, 38 | 56, 62 |

### Statistical analysis

Regressions were performed using stepwise linear regressions (*stepwiselm* function in MatLab’s *Statistics and Machine Learning Toolbox*), so as to select the most parsimonious model. In order to understand what genetic and WM factors are predictive of performance in the Acquire task, we performed a stepwise regression of the seven predictors (three allelic variations, three WM and ethnicity) onto each of several outcome measures representative of performance: success rate, implicit and explicit retention, ΔR, MAD[ΔR], ΔP, MAD[ΔP]. A stepwise logistic regression was employed for overall task success in the Acquire task. For the Preserve task, we performed separate stepwise regressions using the same seven predicators for the following outcome variables: baseline reach direction as a control variable, learning rate in the adaptation block, early and late success rate in the asymptote blocks (first 30 and last 170 trials; Codol et al., 2018), retention in the no-feedback blocks, and ΔR and ΔP during the asymptote blocks.

Prior to the regression analysis, all predictors and predicted variables were standardised (z-scored). For all non-ordinal variables, individual data were considered outliers if further than 3 standard deviations from the mean and were removed prior to standardisation. Multicollinearity of predictors was also assessed before regression with Belsley Collinearity Diagnositcs (*collintest* function in MatLab’s *Econometrics Toolbox*) and no predictors were found to exhibit condition indexes over 30, indicating acceptable levels of collinearity. When considering retention for both tasks, unsuccessful participants were removed from the regression analysis.

In order to quantify the predictive ability of the regression models 10-fold cross validation was performed on the model selected by the stepwise regression. Briefly, this consists of dividing the data samples into 10 evenly sized ‘folds’, the data from nine of the folds are used to create a regression model using ordinary least squares regression and this model is used to predict the values of data in the remaining fold given the values of the predictor variables. We measured the quality of the model fit in the 9 folds (In-sample) and the remaining fold of data (out-of-sample) by calculating the mean absolute error (MAE) of the predicted values from the real values. This process was repeated 1000 times for each model with the data assigned to each fold randomised on every iteration, we present the mean MAE ± SD across the 1000 iterations.

#### Exploratory mediation analysis

We performed a mediation analysis to test if the relationship between SWM and SR was mediated by ΔR. Our hypothesis was that higher SWM enables smaller changes after correct trials (ΔR) and this then explains the relationship between SWM and SR. To ensure that separate trials were used in the calculation of ΔR and SR, we split the trials into two equally sized folds. The SR was then calculated for one fold as a percentage of correct trials, and ΔR was calculated as the median change of reach angle after correct trials in the other fold. For the Acquire task only successful subjects were included in the mediation analysis. We employed Baron & Kenny’s three step mediation analysis (Baron and Kenny, 1986): first regress SR on SWM, then regress ΔR on SWM, and finally regress SR on both SWM and ΔR. Subsequently, we performed a Sobel test to determine if there was a significant reduction in the relationship between SWM and SR when including ΔR. The Sobel test examines if the amount of variance in SR explained by SWM is significantly reduced by including the mediator (Sobel, 1986). For a significant effect to be found, SWM must be a significant predictor of ΔR and ΔR must also be a significant predictor of SR after controlling for the effect of SWM on SR. We repeated this procedure 1000 times with the allocation of trials to each fold randomised on each repetition. We present results in terms of the 95% confidence intervals for the R^2^ values for each of the regressions and the median p-value of the Sobel test, along with the associated 95% confidence intervals.

## Results

### Acquire task

In the Acquire task, participants had to learn to compensate and secretly shifting reward region in order to obtain successful feedback (Figure 2,3). As in Holland et al. (2018), about a quarter (28.1%) of participants failed to learn to compensate for the full extent of the rotation (Figure 3a). Successful participants retained most of the learnt rotation (mean 80.7% ± 28.0% SD) in the maintain block. However, upon being asked to remove any strategy they had been employing, their performance returned to near-baseline levels. Consequently, a large explicit component to retention was found for successful participants (Figure 3b), whereas both successful and unsuccessful participants manifest a small but non-zero implicit component (t(86)=9.90, p=7.43×10^−16^, d=1.061 and t(33)=4.53, p=7.39×10^−5^, d=0.776, respectively; Figure 3c). Furthermore, in accordance with Holland et al (2018), we found that participants made larger (t(120)=15.80, p=4.32×10^−31^, d=1.900) and more variable changes in reach angle following unrewarded trials (t(120)=13.36, p=1.68×10^−25^, d=1.485; Figure 3d-h), whereas in participants who would go on to fail, the post-error adjustments were smaller than in successful participants (t(119)=3.33, p=0.001, d=0.672; Figure 3d). Changes following rewarded trials were similar between the groups (t(119)=0.71, p=0.48, d=0.143; Figure 3f,g). The results obtained in this sample (N=121) therefore replicate results from a previous study (N=30) and provides further confirmation that performance in this task is fundamentally explicitly driven (Holland et al., 2018).

**Figure 2:**
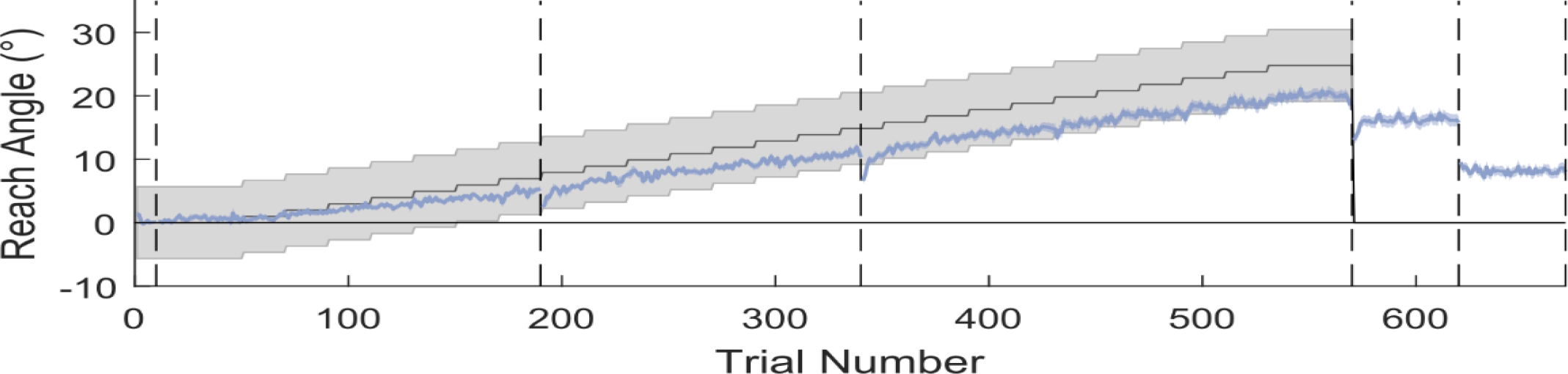
Reaching performance in the Acquire task. The grey region represents the gradually rotating rewarded region, the blue line represents mean reach angle for each trial, and the shaded blue region represent SEM. Vertical dashed lines represent different experiment blocks or breaks. Average performance for the full cohort falls within the reward region and demonstrates a clear reduction in reach angle when asked to remove strategy. N=121.

**Figure 3:**
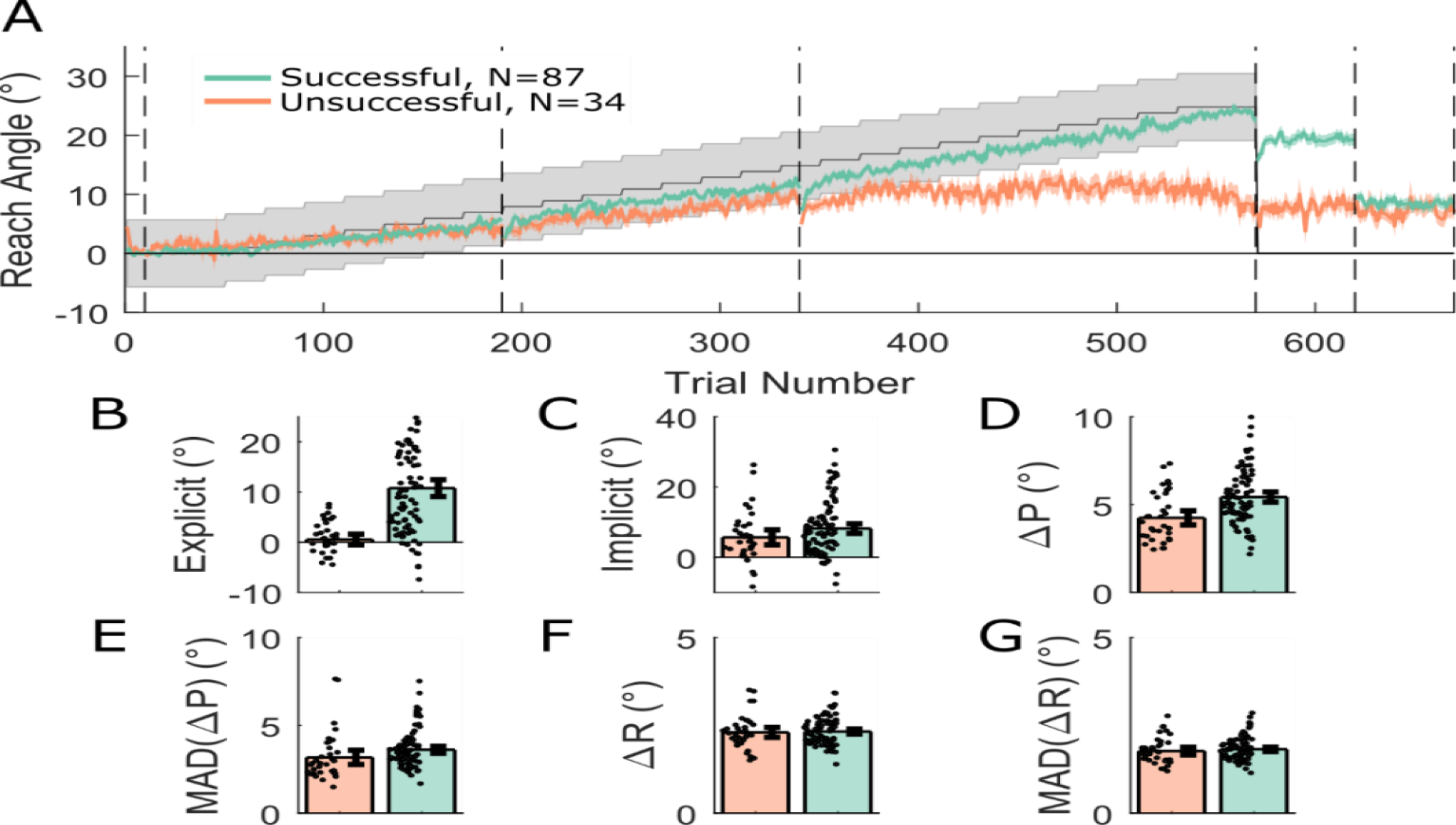
Acquire task split by success at final angle. **A:** Average reach angle for the successful (green) and unsuccessful (orange) groups, shaded regions represent SEM and grey shaded region represents the rewarded region. Despite similar initial performance, a clear divergence can be seen between the two groups and an explicit component to retention is only visible in the successful group, whereas implicit retention is similar between groups. **B-G:** subplots displaying derived measures, which acted as outcome variables for the regression analysis, separated into successful and unsuccessful participants overlaid with individual data points. Error bars represent 95% bootstrapped confidence intervals.

In order to understand what genetic and WM factors are predictive of performance in the reaching task, we performed a stepwise regression of the seven predictors (three allelic variations, three WM and ethnicity) onto each of several outcome measures representative of performance: success rate, implicit and explicit retention, ΔR, MAD[ΔR], ΔP, MAD[ΔP]. Additionally, we performed a stepwise logistic regression of the predictors onto a binary variable encoding if a participant successfully learnt the full rotation (1) or not (0). The logistic regression showed no significant predictors of task success, that is, of being able to follow the shifting reward region until the end of the learning block. However, higher SWM was predictive of an increased success rate (percentage of correct trials; β=0.416, p=2.95×10^−6^). To ensure that the relationship between SWM and success rate was not due to failure at a later point in the task, success rate was measured during the initial period in which subjects who could not fully account for the displacement are still successful; for those who could, the full learning block was included. Next, retention was assessed by splitting up the explicit and implicit components such as in Holland et al. (2018). No predictor was related to the implicit component, but the explicit component was strongly and positively associated with RWM (β=0.373, p=1.78×10^−4^). These results suggest positive relationships for both RWM and SWM with task performance: greater RWM predicts a greater contribution of explicit processes to learning, whereas greater SWM predicts a greater percentage of correct trials.

In Holland et al (2018), the amplitude of the changes in reach angle participants made following unrewarded trials was found to be predictive of task success, that is, greater ΔP was predictive of an increased chance of overall task success. Thus it could be that RWM and SWM, that are shown to associate with performance in this study, are themselves predictors of changes in reach angle. The regression results demonstrated that a large ΔR was inversely related to SWM (β=-0.251, p=0.006), as was MAD[ΔR] (β=-0.283, p=0.002). The results indicate that greater SWM was predictive of smaller and less variable changes in reach angle after successful trials, suggesting high SWM enables the maintenance of rewarding reach angles. It was also found that the variability of changes in reach angle after unrewarded trials (MAD[ΔP]) was negatively predicted by RWM (β=-0.236, p=0.011). This result was unexpected as it suggests that greater WM capacity predicts smaller changes following unrewarded trials, whereas previous results suggest a positive relationship between these changes and overall task success. Finally, to ensure the robustness of the results, we tested whether retaining only the largest ethnic group in our population (i.e. Caucasian, N=82/121) produced the same results as was observed with the full participant pool. In accordance with the full sample, all previously described relationships were also found in the Caucasian only subset (Table 2).

**Table 2:**
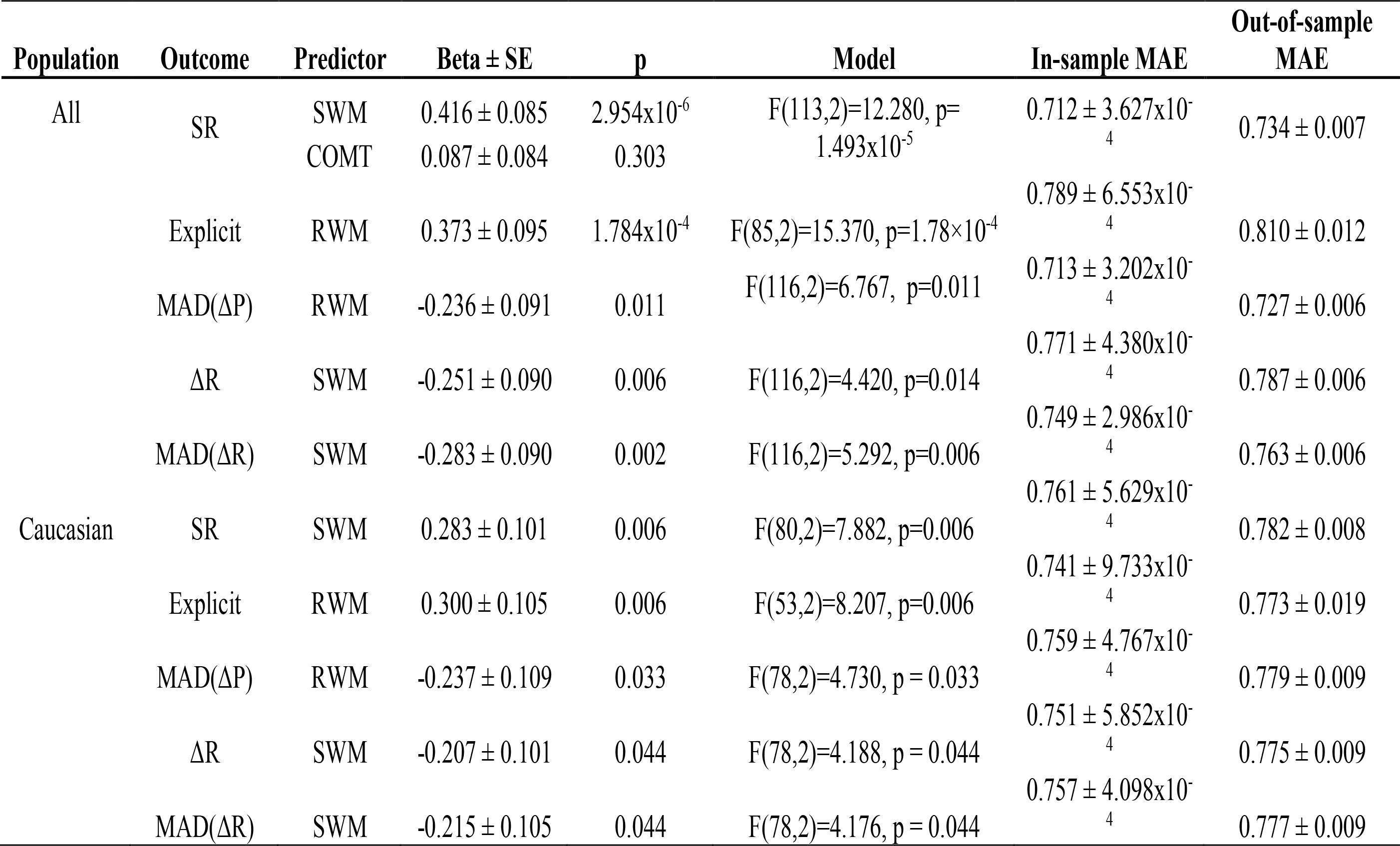
Regressions with significant models for the Acquire task. The predictors selected by the stepwise regression procedure to have a model significantly better than the intercept only model are reported. For each model the selected predictors are reported alongside the coefficient and standard error and associated p value for that predictor, as well as the significance of the model overall. The results of the 10-fold cross-validation analysis are reported in terms of the mean ± SD of the absolute error (MAE) of the model prediction for the 1000 repetitions. Results are reported when including all participants (N=121) or the Caucasian only subset (N=82), demonstrating that the reported results are consistent in both.

Overall, WM (in particular RWM and SWM) successfully predicted various aspects of performance in the Acquire task, while genetic predictors failed to do so. Specifically, greater SWM predicted smaller and less variable changes after correct trials. This suggests that SWM underlies one’s capacity to preserve and consistently express an acquired reach direction to obtain reward. Furthermore, SWM also directly predicted success rate. In addition, greater RWM was a strong predictor of explicit control. The inverse relationship between RWM and the variability of changes after unrewarded trials was unexpected. However, one possible explanation is that participants with poorer WM capacity make larger errors which require larger corrections. Restricting our group to Caucasians showed that these effects are robust to ethnicity.

### Preserve task

In this task, we addressed how well participants can maintain a previously learnt adaptation after transitioning to binary feedback. As participants are unable to compensate for a large abrupt displacement of a hidden reward region (van der Kooij and Overvliet, 2016; Manley et al., 2014), participants first adapted to an abruptly introduced 20° clockwise rotation with full vision of the cursor available. Subsequently, visual feedback of the cursor position was replaced with binary feedback; participants were rewarded if they continued reaching towards the same angle that resulted in the cursor hitting the target during the adaptation phase. Overall, participants adapted to the visuomotor rotation successfully (Figure 4,5a-c) before transitioning to the binary feedback-based asymptote blocks. However, from the start of the asymptote blocks onward, participants exhibited very poor performance, expressing an average 45.0 ± 24.2 SD% success rate when considering all 200 asymptote trials (Figure 4,5a, d,e). Separating successful and unsuccessful participants (40% success rate cut-off; Figure 5a) revealed that successful participants expressed behaviour greatly similar to that observed in Codol et al. (2018), in which unsuccessful participants were excluded, using the same cut-off (40% success rate). The ‘spiking’ behaviour observed in reach angles during the asymptote blocks (Figure 5a) is due to the presence of the ‘refresher’ trials, with the large positive changes in reach angle corresponding to trials immediately following the refresher trials. This pattern of behaviour is particularly pronounced in the unsuccessful participants. Finally, participants demonstrated at least a residual level of retention even after being instructed to remove any strategy they had employed (t(69)=7.268, p=3.345×10^−10^, d=0.869; Figure 5a,f). Therefore, the results obtained in this sample (N=120) replicate results from a previous study (Codol et al., 2018; N=20, BF-Remove group) and provides further confirmation that performance in this task is fundamentally explicitly driven. As with the Acquire task successful participants displayed larger changes in angle after unrewarded trials than their unsuccessful counterparts (t(117)=3.847, p=1.952×10^−4^, d=0.717; Figure 5h). However, in contrast to the Acquire task, successful participants also displayed smaller changes in angle after rewarded trials (t(115)=-7.534, p=1.218×10^−11^, d=1.421; Figure 5g).

**Figure 4:**
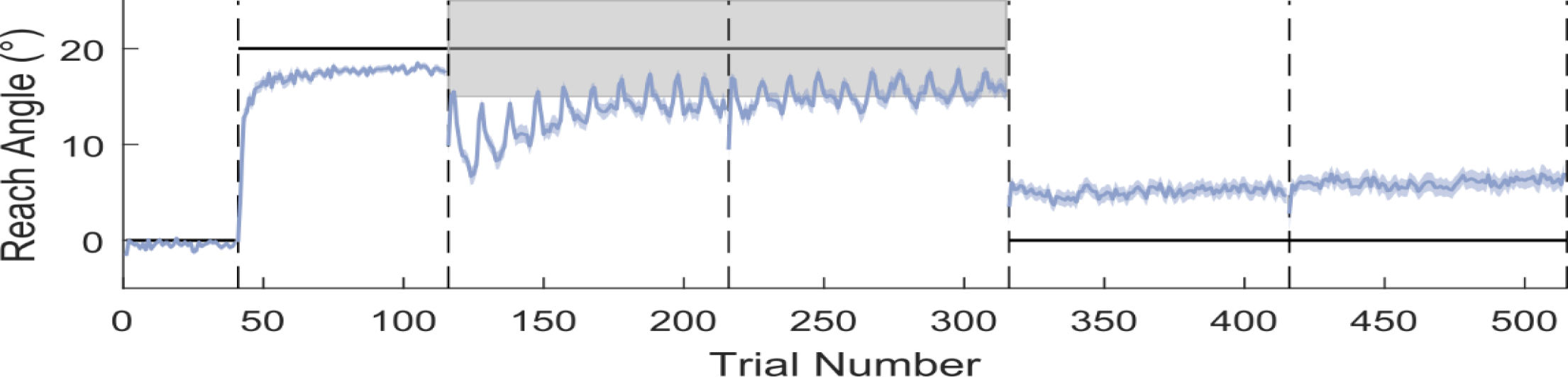
Reaching performance in the Preserve task. The grey shaded area represents the rewarded region, and the thick black line represents the perturbation. The vertical dashed lines represent block limits. The blue line indicates mean reach angle for every trial and blue shaded areas represent SEM. After successfully adapting to the visuomotor rotation performance deteriorates at the onset of binary feedback, subsequently success rate increases towards the end of the asymptote blocks. Following the removal of all feedback, and the instruction to remove any strategy, a small amount of implicit retention remains. N=120.

**Figure 5:**
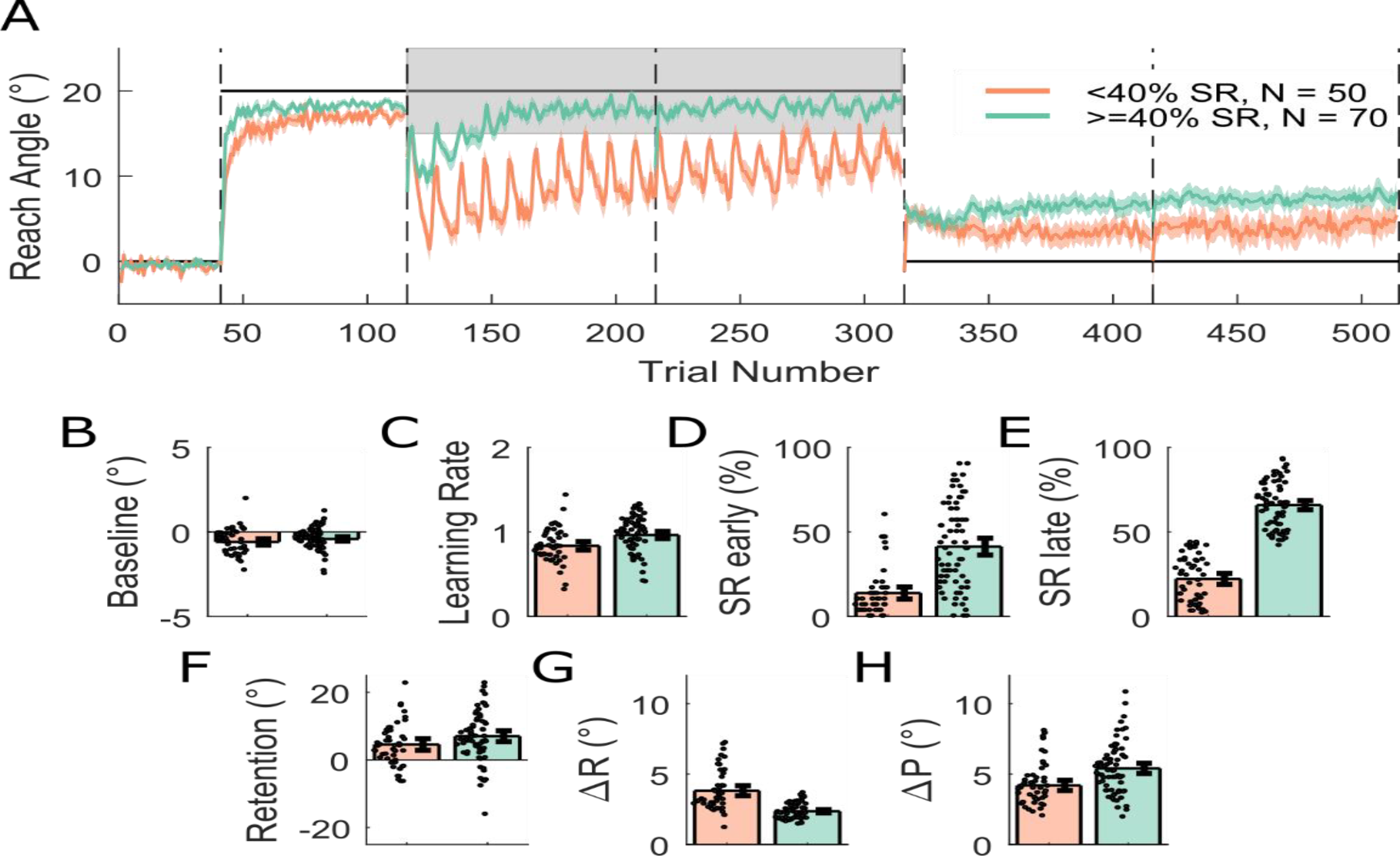
Preserve task split into two groups on the basis of success rate. **A:** Shaded regions represent SEM. **B-H:** Derived variables, which acted as outcome variables for the regression analysis, for the two groups, error bars on the bars represent 95% bootstrapped confidence intervals and individual data points are displayed. SR: success rate.

As in the Acquire task, we examined if performance in any of the WM tasks or genetic profile could predict participant’s behaviour in the reaching task. We performed separate stepwise regressions for the following outcome variables: baseline reach direction as a control variable, learning rate in the adaptation block, early and late success rate in the asymptote blocks (first 30 and last 170 trials; Codol et al., 2018), retention in the no-feedback blocks, and ΔR and ΔP during the asymptote blocks. The most striking result was that both early and late success rate could be reliably predicted by RWM (early: β=0.255, p=0.005; late: β=0.287, p=0.002; Table 3), with greater RWM associated with increased success rates.

**Table 3.**
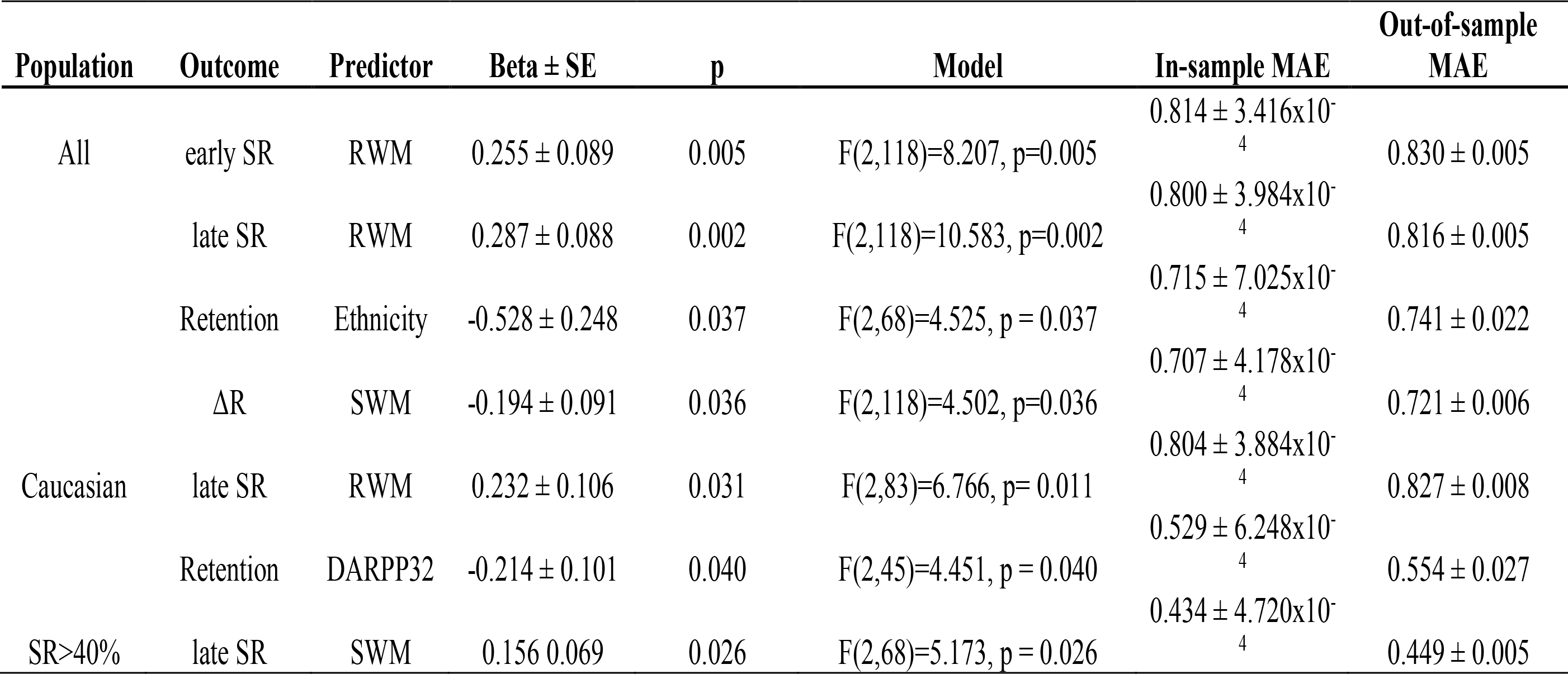
Regression with significant models for Preserve task. The predictors selected by the stepwise regression procedure to have a model significantly better than the intercept only model are reported. For each model the selected predictors are reported alongside the coefficient and standard error and associated p value for that predictor, as well as the significance of the model overall. The results of the 10-fold cross-validation analysis are reported in terms of the mean ± SD of the absolute error (MAE) of the model prediction for the 1000 repetitions. Results are reported when including all participants (N=120) or the Caucasian only subset (N=85), demonstrating that the relationship between RWM and late success rate are consistent in both and revealing a genetic predictor of retention.

Genetic profile did not predict any aspect of performance, analogous to the Acquire task. In contrast, greater SWM successfully predicted reduced ΔR (β=-0.194, p=0.036), similarly to the Acquire task. Finally, retention values were surprisingly predicted by ethnicity (β=-0.528, p=0.037). Due to the existence of a relationship between ethnicity and retention, we performed the same analysis as in the Acquire task, that is, we tested if our observed results hold if only our largest ethnic group (Caucasian, N=85/120) was considered. In accordance with the result for the full population, the positive relationship between late success rate and RWM was again observed (β=0.232, p=0.031). However, the SWM-ΔR and RWM-early success rate relationships were no longer observed in this smaller subset of the population. Interestingly, retention was now predicted by a genetic variable, DARPP32 (β=-0.214, p=0.040), suggesting that less dopaminergic metabolism leads to better retention. This last result again suggests a possible confound, that is, that DARPP32 allelic distribution was different across ethnic groups. However, a χ^2^ test analysis demonstrated that DARPP32 alleles were evenly distributed between the Caucasian and non-Caucasian group, ruling out this possibility (χ^2^=2.578, p=0.276). As a post-hoc analysis we performed the same stepwise regressions for the outcome variables success rate early and success rate late but restricted to participants with an overall success rate of greater than 40%. Although we found no predictors of early success rate, we did find that higher SWM was predictive of a higher late success rate (β=0.156, p=0.026). This result is in contrast to the relationship of RWM to late success rate when including all participants.

Overall the regression results fit a pattern similar to that found for the Acquire task with greater RWM predicting improved performance on the reaching task and greater SWM predicting smaller changes in reach angle after rewarded trials. However, in the Preserve task in one specific instance we did observe a genetic predictor of performance.

### Cross validation analysis

To test the predictive ability of the regression models we performed 10-fold cross validation on the final model selected by the stepwise regression process. The quality of the in-sample and out-of-sample fits was assessed by calculating the MAE. From Tables 2 and 3 it can be seen that although the out-of-sample MAE is consistently higher than that of the corresponding in-sample, all differences are less than 0.1 and all of out-of-sample MAEs are below 1. As both the predictor and outcome variables are standardised this indicates that the mean error of the prediction was less than 1 standard deviation of the outcome variable, and the small increases observed between the in-sample and out-of-sample indicates that the models make accurate predictions when confronted with data on which they were not trained.

### Exploratory analysis

As a relationship exists between SWM and ΔR in both the Acquire and Preserve paradigms, we ran exploratory regressions to assess the relationship between ΔR and success rate across both tasks. Since ΔR and success rate are conceptually strongly related variables, and measuring on the same data set would render them non-independent, we split each individual’s reaching data into two sections and assessed whether ΔR or ΔP in the first section could reliably predict success rate in the second (see methods for details). Although we found no predictors of ΔP in our primary analysis, results here in combination with previous work (Holland et al., 2018) has demonstrated a link between ΔP and task success, with a greater ΔP indicative of greater success. Therefore, we also performed the same analysis for ΔP.

In the Acquire task, ΔR and ΔP in the first section of learning trials predicted success rate in the final twenty trials, though ΔP appeared as the strongest predictor (ΔR: β=-0.274, p=0.015; ΔP: β=0.581, p=3.89×10^−6^; Figure 6a,b, yellow; Table 4). Similarly, for the Preserve task ΔR and ΔP in the first half of asymptote trials predicted success rate in the second half (ΔR: β=-0.750, p=1.07×10^−12^; ΔP: β=0.229, p=0.007; Figure 6a,b, pink; Table 4). In both tasks, the directions of these relationships were opposite; greater success rate was predicted by smaller changes following rewarded trials and greater changes following unrewarded trials. In summary, we found that for both tasks the magnitude of changes in behaviour in response to rewarded and unrewarded trials early in learning were strongly predictive of future task success across both the Acquire and Preserve tasks.

**Table 4:** Regression results for split data for both the Acquire and Preserve tasks. Ordinary least squares linear regressions were performed with both ΔR and ΔP included as predictors. The regression coefficient, standard error and p value for each predictor are reported along with the significance of the comparison between the model and an intercept only model. In both tasks there is an opposing relationship between ΔR and ΔP and success rate, with smaller changes after rewarded trials and larger changes after unrewarded trials predictive of success.

| | | $\Delta R$ | $\Delta P$ | Model |
| --- | --- | --- | --- | --- |
| Acquire | $\beta$ | -0.274 | 0.581 | F(115,2)=11.9<br>p=2.09$\times 10^{-5}$ |
|  | SE | 0.111 | 0.120 |  |
| | p | 0.015 | 3.89$\times 10^{-6}$ | |
| Preserve | $\beta$ | -0.750 | 0.229 | F(112,2)=35.3<br>p=1.28$\times 10^{-12}$ |
|  | SE | 0.093 | 0.084 |  |
| | p | 1.07$\times 10^{-12}$ | 0.007 | |

**Figure 6:**
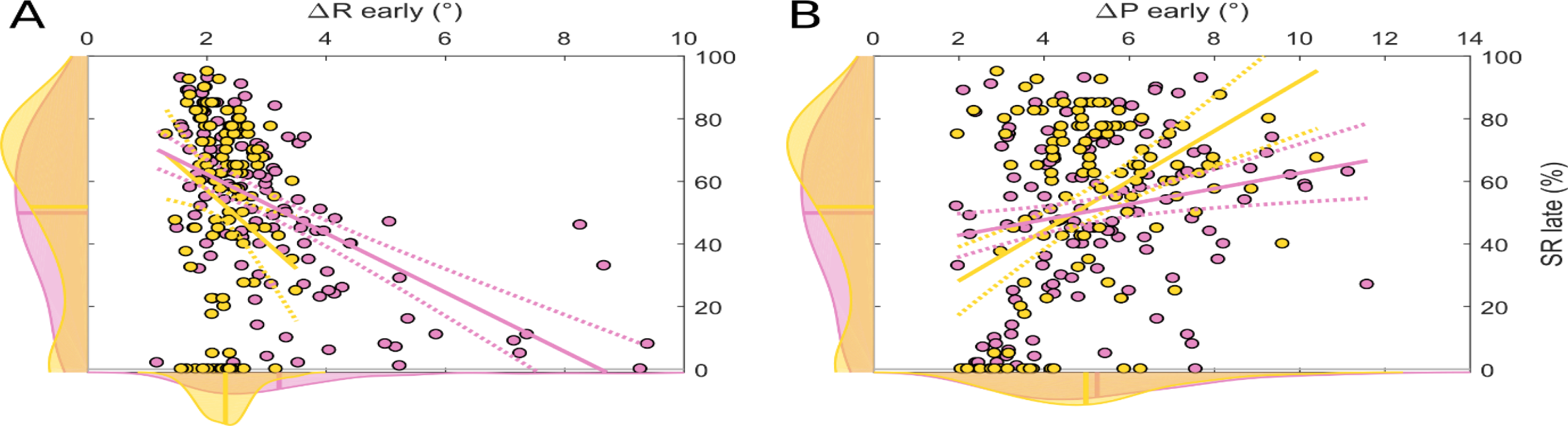
Slice plots showing regression results for prediction of late success rate (SR) by changes in reach angle following rewarded (A) and unrewarded (B) trials during the early learning period. The central axis of each panel displays the individual data from the Acquire (yellow) and Preserve (pink) task, the smoothed distribution of the data in each dimension is displayed on the corresponding axis. Solid lines represent the prediction of the regression model when the other predictor is held at its mean value.

**Figure 7.**
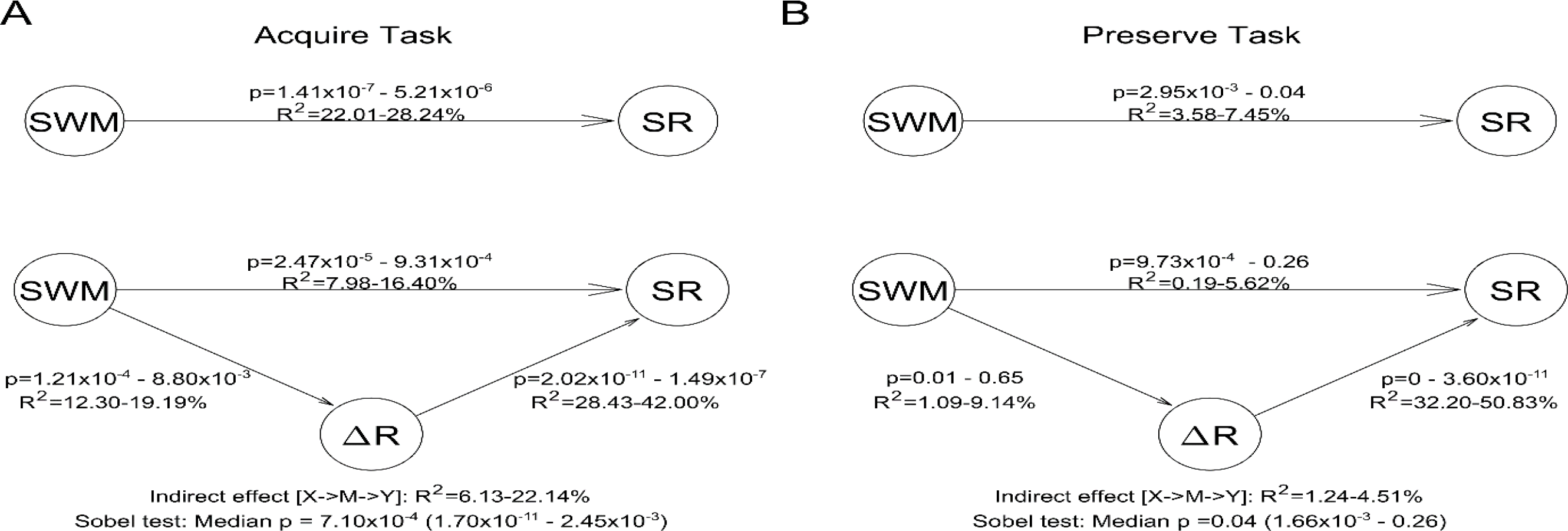
Mediation Analysis for both the Acquire (A) and Preserve (B) tasks. The numbers associated with each arrow display the 95% confidence intervals for each of the relationships (R^2^ and p-values) across the 1000 repetitions. Below the figure, the results of the Sobel test are displayed indicating the amount of variance explained by the indirect pathway and the 95% confidence intervals and median p-value.

### Mediation analysis

To test whether the effect observed between SWM and SR was explained by an indirect effect through ΔR, we performed an exploratory mediation analysis on both tasks. For both the Acquire and Preserve tasks, the results indicate a significant proportion (median p=7.10×10^−4^ and p=0.04 respectively) of the relationship between SWM and SR can be explained by a mediation from SWM via ΔR to SR. However, in the case of the Acquire task, a significant relationship between SWM and SR still exists, indicating that not all of the effect of SWM on SR could be explained by the indirect pathway. Whereas, in the Preserve task the SWM-ΔR relationship is weaker and was not significant on every repetition, occasionally leading to an insignificant mediation effect, despite the median p-value indicating an effect when considering all repetitions.

## Discussion

In this study, we sought to identify if genetic background or specific domains of WMC could explain the variability observed in performance levels during reward-based motor learning tasks. We found that RWM and SWM predicted different aspects of the Acquire and Preserve tasks, whereas VWM did not relate to any performance measure. Specifically, RWM predicted the explicit component of retention in the Acquire task and success rate in the Preserve task, whereas SWM predicted success rate in the Acquire task and ΔR in both tasks. Conversely, allelic variations of the three dopamine-related genes (DRD2, COMT and DARPP32) did not predict any behavioural variables in the full sample of participants. This suggests that SWM predicts a participant’s capacity to reproduce a rewarded motor action, while RWM predicts a participant’s ability to express an explicit strategy when making large behavioural adjustments. Therefore, we conclude that WMC plays a pivotal role in determining individual ability in reward-based motor learning.

Recently, Wong et al. (2019) describe a positive relationship between SWM and the development of explicit strategies in visuomotor adaptation, complimenting previous reports (Anguera et al., 2012; Christou et al., 2016). However, in contrast to the current findings the previous experiments employed relatively small sample sizes, which may render correlations unreliable. The large group sizes employed here and the confirmation of relationships across two tasks provides strong evidence that these relationships are robust, replicable, and extend from visuomotor adaptation to reward-based motor learning. An interesting dichotomy was the reliance on SWM and RWM for the Acquire and Preserve task, respectively. The Preserve task required the maintenance of a large, abrupt behavioural change, whereas the Acquire task required the gradual adjustment of behaviour considering the outcomes of recent trials. Therefore, RWM may underscore one’s capacity to express a large correction consistently over trials with binary feedback, whereas SWM reflects one’s capacity to maintain a memory of previously rewarded actions and adjust behaviour accordingly. Conformingly, the magnitude of ΔR was negatively related to SWM but not RWM in both tasks, suggesting high SWM enables the maintenance of rewarding actions. Supporting this, Sidarta et al. (2018) report that higher SWM associated with reduced movement variability in a reward-based motor learning task. Additionally, explicit retention, an element of the Acquire task requiring a large, sudden change in reach direction, was predicted not by RWM not SWM.

A notable feature of the Preserve task is the “spiking” behaviour observed immediately following ‘refresher’ trials, suggesting a central role of ‘refresher’ trials in binary feedback-based performance when included (Codol et al., 2018; Shmuelof et al., 2012). The transient nature of this decrease in error demonstrates this is insufficient to promote generalisation to binary feedback trials, at least in unsuccessful participants. It remains an open question whether superior performance of successful participants was partly due to a capacity to generalise information from ‘refresher’ trials. McDougle and Taylor (2019) provided evidence that two separate strategies are employed in visuomotor adaptation; response-caching and mental rotation. The balance between the two strategies is a function of task demands. It could be that the relationships between RWM and SWM to success rate in the Preserve and Acquire tasks respectively reflect a different balance of the use of these strategies; visual feedback in ‘refresher’ trials in the Preserve task encourages the engagement of mental rotation processes, whereas the slow updating of behaviour in the Acquire task in the absence of visual feedback engages the response-caching memory system. This would imply that response-caching is associated with SWM.

Surprisingly, although ΔP was a strong predictor of success in both tasks, it was not predicted by any genetic variable. In the Acquire task MAD(ΔP) was inversely predicted by RWM. This result is surprising given the positive relationship between ΔP and success rate in both tasks. Whilst no predictor of ΔP was found in the Preserve task, ΔP is however likely important for explicit control, as errors are a central element leading to the induction of structural learning in reward-based tasks, reinforcement learning (Daw et al., 2011; Manley et al., 2014; Sutton and Barto, 1998) and motor learning in general (Maxwell et al., 2001; Sidarta et al., 2018).

If RWM is important for explicit control and the main element predicting success in the Preserve task, this raises the question as to whether gradual designs (as in the Acquire task) are more suitable to engage implicit reinforcement learning, at least initially. However, the Acquire task still bears a strong explicit component (Holland et al., 2018). How can these two views be reconciled? In reward-based motor learning tasks, it is generally agreed that participants begin to reflect upon task structure and develop strategies upon encountering negative outcomes (Leow et al., 2016; Loonis et al., 2017; Manley et al., 2014; Maxwell et al., 2001), which occurs nearly immediately in the Preserve task after the introduction of binary feedback, due to a lack of generalisation of cerebellar memory (Codol et al., 2018). In contrast, in the Acquire task, participants experience an early learning phase with mainly rewarding outcomes, possibly suppressing development of explicit control and allowing for this early window of implicit reward-based learning. Other studies have demonstrated that minor adjustments in reach direction under reward-based feedback can occur, though none has assessed their explicitness directly in the very early stages, such as about 1-4° (Izawa and Shadmehr, 2011; Pekny et al., 2015; Therrien et al., 2016). Notably, Izawa and Shadmehr, (2011) observed that after 8° shifts of a similarly-sized reward region, participants indeed noticed the perturbation, but awareness was not assessed for smaller shifts.

In Holland et al., (2018), the addition of a RWM-like dual-task was very effective in preventing explicit control, leading to participants invariably failing at the reaching task. It may therefore appear as surprising that RWM does not predict success rate in the Acquire task. A possible explanation is that RWM and SWM share the same memory buffer (Anguera et al., 2010; Beschin et al., 2005; Cohen et al., 1996; Jordan et al., 2001; Suchan et al., 2006). Similarly, in force-field adaptation the early component of adaptation – considered as bearing a strong explicit element – is selectively disrupted with a VWM dual-task (Keisler and Shadmehr, 2010). However, we found no relationships with VWM in our reward-based motor tasks. It may be possible that reward-based motor performance relies more on spatial instances of WM as opposed to tasks such as force-field adaptation.

The absence of DA-related genetic relationships with behaviour is a surprising result as a substantial body of literature points to a relationship between dopamine and performance in reward-based tasks, including those with motor components (Deserno et al., 2015; Doll et al., 2016; Frank et al., 2007, 2009; Gershman and Schoenbaum, 2017; Izawa and Shadmehr, 2011; Nakahara and Hikosaka, 2012; Pekny et al., 2015; Therrien et al., 2016). There exists a growing appreciation of the links between decision-making and motor learning (Chen et al., 2018; Haith and Krakauer, 2013). Chen et al., (2017) demonstrated that exploratory motor learning can be modelled as a sequential decision-making process, with individual’s explorative drive shared between motor and decision-making tasks. However, the results presented here suggest that genetic predictors of exploration and exploitation in decision-making tasks are not also predictive of similar behaviours in reward-based motor learning.

Our sample sizes were defined *a priori* for 90% power based on previous work (Doll et al., 2016; Frank et al., 2009; see pre-registrations), therefore they are unlikely to be underpowered. Another possibility is that we employed the wrong variables to assess behaviour. However, given the informative and coherent relationships between WM and motor learning and the ability to predict overall performance on that basis, could it be that the genes we selected do not relate in any meaningful way to performance in these reward-based tasks? In line with this, a recent study showed that DA pharmacological manipulation did not alter reward effects in a visuomotor adaptation task (Quattrocchi et al., 2018). However, previous work has shown that Parkinson’s disease patients show impaired reward-based motor performance (Pekny et al., 2015). It is possible that genetic variations may not impact reward-based motor learning significantly, especially compared to the wide depletion of dopaminergic neurons in Parkinson’s disease. Finally, future work should also address the possible involvement of other neuromodulators, such as acetylcholine, norepinephrine and serotonin (for a review, see Dash et al., 2007), in reward-based motor learning.

In summary, despite employing two distinct tasks and an independent participant pool on different devices, we find strikingly similar results in reward-based motor learning. While SWM strongly predicted a participant’s capacity to reproduce successful motor actions, RWM predicted a participant’s ability to express an explicit strategy when required to make large behavioural adjustments. Therefore, both SWM and RWM are reliable predictors of success during reward-based motor learning. Surprisingly, no dopamine-related genotypes predicted performance. Therefore, WMC plays a pivotal in determining individual ability in reward-based motor learning. This could have important implications when using reward-based feedback in applied settings as only a subset of the population may benefit.

## Acknowledgements

This work was supported by the European Research Council starting grant: MotMotLearn (637488)

